# Expanding the Genome in a Bottle Truth Set: Detection and Validation of Novel Low-frequency Variants Using High-accuracy NanoSeq

**DOI:** 10.64898/2025.12.05.692678

**Authors:** Yang Zhang, Hsu Chao, Muchun Niu, Christopher M. Grochowski, Kavya Kottapalli, Sravya V. Bhamidipati, Donna M. Muzny, Richard A. Gibbs, Chenghang Zong, Harsha Doddapaneni

## Abstract

**Highlights:**

- NanoSeq-MBN achieves near-genome, Poisson-like coverage with minimal trinucleotide bias.
- Expands the GIAB truth set by up to 160k de novo variants.
- Adds a somatic layer to GIAB, enabling benchmarking and calibration of rare variants.
- High-CADD exonic and splice variants highlight value for surveillance and clinical triage.

Somatic mutations record tissue molecular history and inform risk, prognosis, and therapy, yet their variant allele fractions often fall below the reliable detection limit of conventional short-read sequencing. In contrast, duplex sequencing technology featured by NanoSeq applies the principle of single molecule detection and thereby overcomes the limitation. However, the original NanoSeq protocol relies on the restriction enzyme-based genome fragmentation, which constrained its genome coverage to 30-40%. To enable whole-genome discovery with duplex-level fidelity, we pursued two complementary approaches to optimize the NanoSeq protocol: (i) a restriction-enzyme strategy densifies accessible sites using orthogonal 4-bp cutters; and (ii) a workflow using sonication followed by mung bean nuclease with T4 polynucleotide kinase, Klenow fragment and dATP/ddBTP mixture (NanoSeq-MBN) to blunt and repair/A-tailing DNA, while minimizing repair artifacts. We systematically benchmarked their performance using Genome in a Bottle (GIAB) gold-standard sample mixtures. As a result, NanoSeq-MBN achieved near genome-wide, Poisson-like coverage with minimal trinucleotide-context bias and ultra-high accuracy. Beyond variants already present in the GIAB truth set, NanoSeq-MBN identified approximately 120,000-160,000 de novo mutations per sample missing in the truth set, Notably, over 98% had orthogonal support in reanalyzed GIAB bulk Illumina HiSeq libraries. These novel variants extended GIAB from germline benchmarking to rare-variant discovery and calibration of subclonal detection. Functional annotation revealed enrichment of high Combined Annotation Dependent Depletion (CADD) scores mutations in exonic and splice-related regions. Variants intersecting ClinVar entries and OMIM genes highlighted potential for surveillance and clinical triage. Collectively, these results add a somatic layer to GIAB, enabling calibration of burdens and mutational signatures in lymphoblastoid lines and provide reference material for rare-variant assays. The NanoSeq-MBN workflow offers a path to whole-genome, high-fidelity discovery of ultra-rare somatic variation with relevance to clinical assay validation.

## INTRODUCTION

Somatic mosaicism arises because tissues follow distinct developmental trajectories, experience diverse environmental exposures, and endure heterogeneous physiological stresses. Over time these forces produce distinct “molecular history” at different regions of the tissues. Although most somatic variants are neutral, a clinically relevant subset perturbs regulatory or coding sequences and can influence disease risk, prognosis, or therapy—particularly in aging-related diseases, neurodegenerative disorders, and cancer ^1–4^. Accurate discovery of such variants therefore has immediate translational value.

Conventional bulk short-read sequencing has a per-base error rate of ∼10⁻³, restricting confident detection to variants with variant allele frequency (VAF) above 1%—typically early embryonic or clonally expanded events^5^. By contrast, somatic variants in normal tissues^6,7^ often occur at VAFs on the order of 10^-6^, well below this methodological limit. Duplex sequencing overcomes this by independently tagging and comparing both strands, reducing background errors to ≤10-⁷–10⁻⁹ per base and enabling confident detection of ultra-rare mutations^8–11^. NanoSeq^7^, a restriction-enzyme–based duplex strategy, present the lowest error rates of <5×10^-9^ per base. Nevertheless, its restriction-enzyme implementation, samples only ∼30–40% of the genome, limiting discovery—especially in non-coding regions distant from enzyme cut sites^12^.

To overcome this coverage constraint while preserving fidelity, we pursued two complementary optimizations within the NanoSeq framework: (i) increasing restrictionenzyme (RE) diversity (adding *AluI* and *Hpy166II* to HpyCH4V) to maximize accessible cut sites across the genome, and (ii) enabling enzyme-agnostic fragmentation via sonication, followed by blunt-end repair using mung bean nuclease (MBN) optionally supplemented with T4 polynucleotide kinase (T4PNK), end repair and Klenow fragment (3’ -> 5’ exo-)/dATP/ddBTP for DNA repair and A-tailing^13^ to minimize polymerase-induced artifacts. This strategy aims for near-whole-genome, uniform coverage without sacrificing duplex-level accuracy.

Taking advantage of Genome-in-a-Bottle (GIAB) gold-standard samples—HG001 (NA12878, European ancestry), HG002 (Ashkenazi Jewish ancestry), and HG005 (Han Chinese ancestry) ^14–16^, we experimentally generated a GIAB sample mixture so that we have groundtruth information of low VAF mutations (REF). We optimized and systematically benchmarked different NanoSeq protocols with the GIAB sample mixture, as well as the low-mutation-burden control of a human cord blood sample. As a result, we not only validated the different NanoSeq protocols, but also uncovered novel low-frequency variants previously not covered by the GIAB truthset. Such information will provide a critical foundation for validating^16^.

## RESULTS

### Complementary designs extend accessible genome while preserving duplex fidelity

Two complementary strategies were implemented to expand the fraction of genome amenable to sequencing without inflating errors. First, RE fragmentation density was increased by sequentially combining enzymes with orthogonal 4-bp recognition motifs (HpyCH4V → HpyCH4V+AluI → HpyCH4V+AluI+Hpy166II). Second, we adopted an enzyme-agnostic fragmentation workflow involving sonication followed by blunt-end repair using MBN, optionally supplemented with T4PNK to restore 5′-phosphates and enhance adapter ligation efficiency. Together, these five library designs—three REbased and two MBN-based (with or without T4PNK)—allow a direct comparison between concentrated power around restriction sites and broad, near–whole-genome accessibility. Library quality control metrics are provided in Table S1.

To quantify detection sensitivity and accuracy under controlled yet genome-wide conditions, a GIAB-based spike-in mixture was prepared by combining genomic DNA from HG005, HG001, and HG002 in a 60:2:1 mass ratio (qPCR-verified; Methods), hereafter referred to as the three-sample mix. Leveraging the high-confidence genotypes for each donor, the expected VAF at every callable locus was derived as the copy-number–weighted average of allele dosage across donors.

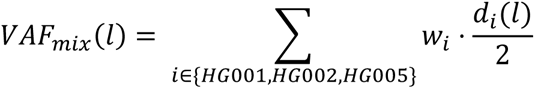

where *w*_*i*_ is the donor’s mass fraction (normalized to 1) and *d*_*i*_(*l*) ∈ {0,1,2} is donorspecific alternate-allele dosage at locus *i* (for autosomes; sex chromosomes handled analogously). Aggregating over loci, mutations were classified into 26 mutually exclusive genotype classes that encode the presence/absence and zygosity pattern across the three donors (3³ combinations minus the all-reference state), yielding a theoretically predicted VAF distribution per class (Figure 1A, *l*). Given a library’s duplex depth at *l* and its consensus criteria, the class-specific probability of detection follows directly from a binomial sampling model. This calibration frames downstream analyses in terms of expected versus observed detection at the level of genotype classes rather than raw counts, thereby controlling mixture composition, allelic dosage, and ploidy.

**Figure 1.**
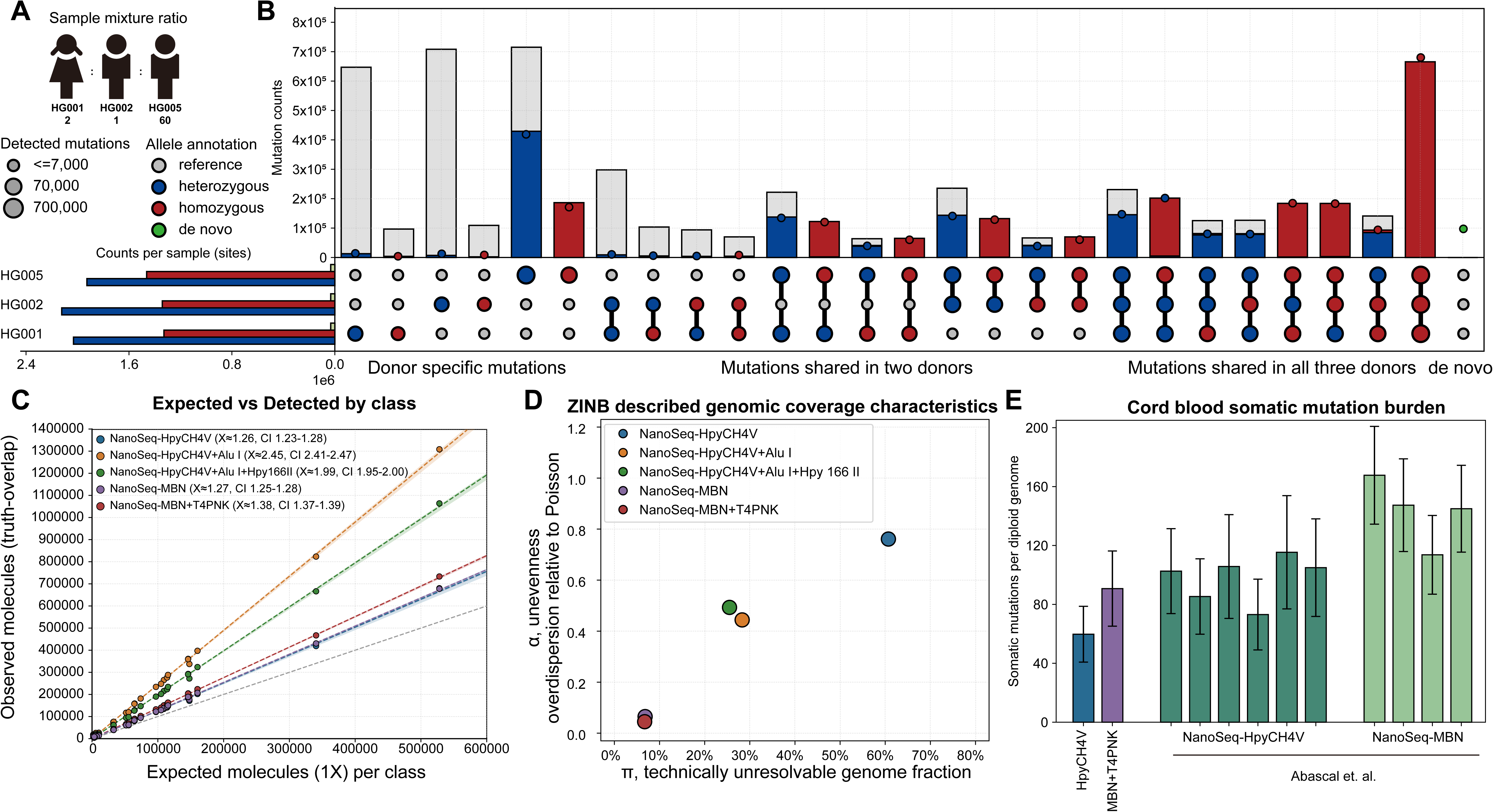
Benchmarking five NanoSeq strategies. A) Sample design: HG001, HG002, and HG005 mixed at a 2:1:60 DNA-mass ratio. B) Target-variant stratification in the GIAB three sample-mixture: lower-left, total germline mutations per individual (histogram); lower-right, genotype combinations across the trio (dot plot); upper-right, stacked bars showing total mutations per genotype (gray) and expected molecule-detection contributions from each donor (colors). C) Expected vs. observed molecule detections by class; linear fits approach unity across libraries. D) Genome coverage breadth and evenness from ZINB modeling; both NanoSeq-MBN libraries show unbiased whole-genome coverage. E) Cordblood somatic mutation burden; NanoSeq-MBN is comparable to the original restriction-based NanoSeq.

We next applied the above framework to the five library designs. For each design origin-constrained regressions were fitted between the expected and observed counts of unique duplex molecules carrying each mutation across 26 variant classes (Figure 1C, Table S2). The regression slopes provide an interpretable measure of effective molecule depth, whereas deviations from linearity reflect class-specific bias. All five libraries exhibited near-perfect linearity (R² = 0.9985–0.9992), confirming that mutation recovery scales proportionally with theoretical expectation. These results demonstrate chemistry-agnostic fidelity of duplex consensus and validating the use of class-aware calibration for downstream comparisons.

### Coverage breadth and evenness favor MBN designs

To assess genomic accessibility and coverage uniformity, locus-level duplex molecule counts were modelled using a zero-inflated negative-binomial (ZINB) distribution (Figure S2, Method). The model partitions observed coverage zeros into two components: the zero-inflation parameter (π), representing the fraction of loci that are technically unresolvable regardless of sequencing depth, and the overdispersion parameter (α), capturing stochastic variability in coverage beyond Poisson expectation.

Among RE designs, the single-enzyme NanoSeq-HpyCH4V library exhibited a high zero-inflation parameter (π ≈ 0.6), indicating that approximately 60% of the genome remained technically inaccessible (Figure 1D). Incorporating *AluI* alongside HpyCH4V increased the resolvable fraction to ∼70%, while further addition of Hpy166II provided only a modest ∼5% gain—likely because excessive fragmentation shortened a subset of molecules below the library capture threshold. Two- and three-enzyme combinations also reduced over-dispersion approximately two-fold relative to the single-enzyme design, reflecting improved coverage evenness. In contrast, both NanoSeq-MBN libraries achieved near-complete genome coverage (>93%) with coverage distributions approaching Poisson expectation (α approaches zero), indicating minimal locus-to-locus variability. Collectively, these results show that while multi-enzyme RE strategies modestly expand accessible regions and improve uniformity, their performance plateaus, whereas the MBN workflow achieves a qualitative improvement in both breadth and evenness unattainable by restriction-based designs.

### *De novo* variants

In addition to the designed germline variants, approximately one hundred thousand *de novo* variants absent from the GIAB reference set were detected across multiple libraries (Figure S1). To assess whether these represented genuine biological variants or technical artifacts introduced during library preparation, the original NanoSeq and NanoSeq-MBN workflows were benchmarked on a low mutation burden cord blood control. As shown in Figure 1E, the NanoSeq-MBN with T4PNK design exhibited only a marginal inflation of apparent somatic burden (Fisher’s exact test *p = 0.0524*), consistent with the fidelity previously reported^7^. These results demonstrate that NanoSeq-MBN with T4PNK achieves near–whole-genome coverage while maintaining Duplex-seq accuracy, establishing a reliable baseline for subsequent variant interpretation.

### Novel variant(s) classification

Building upon this specificity benchmark, single-sample duplex sequencing with MBN+T4PNK was conducted (Table S3) to investigate the origin of the observed de novo variants. To exclude the possibility that such variants arose during cell propagation after the GIAB datasets were established, the original bulk HiSeq data for HG001, HG002, and HG005 were reanalyzed after down-sampling to 30x coverage and alignment to GRCh38. Across all three individuals, approximately 100,000 variants exhibited orthogonal support in the bulk datasets (Figure 2A), validating them as bona fide genomic variants and extending the GIAB truth resource.

**Figure 2.**
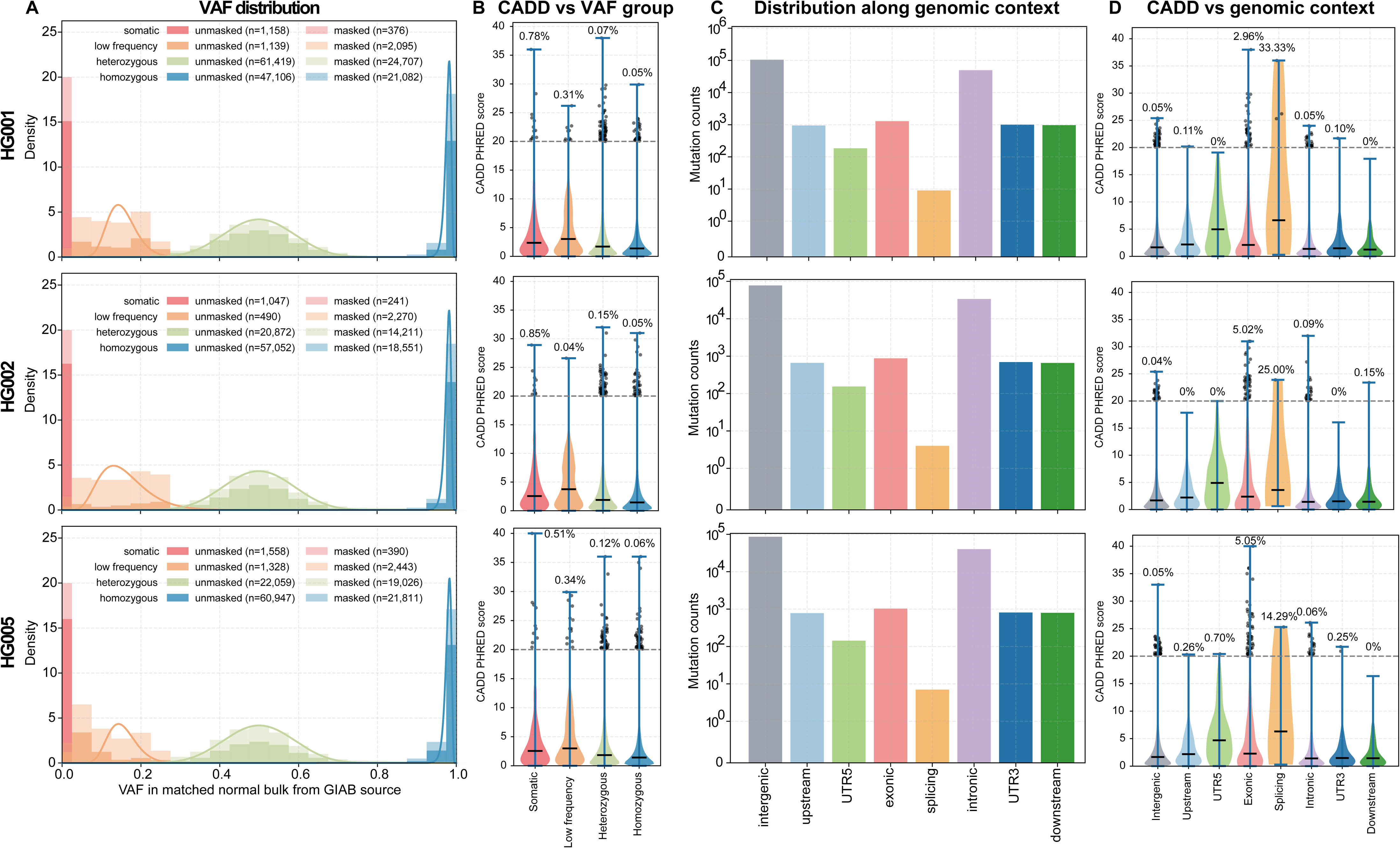
de novo mutation stratification. Rows: the three GIAB individuals (HG001, HG002, HG005). Columns: A) VAF distribution for de novo mutations. B) CADD score by VAF group (violin). C) Genomic context distribution (bar): intergenic, intronic, exonic, UTR, promoter, etc. D) CADD score by genomic context (violin).

To resolve the allelic composition and potential origins of *de novo* variants, a logit-Gaussian mixture model (logit-GMM) was applied to VAF distribution. This framework classifies these *de novo* variants into four categories—homozygous-like, heterozygous-like, low-frequency, and somatic—based on mixture component assignments. Somatic variants were defined as variants with sufficient read depth in the bulk dataset but no alternative alleles. Across samples, the majority of *de novo* variants clustered at VAFs near 0.5 or 1.0, consistent with missed germline mutations that were excluded during the stringent multi-center curation of the GIAB resource. In addition, approximately 3,000 low-frequency variants (VAF <0.3) and 1,500 somatic variants were identified per sample, indicating the presence of subclonal or even single cell private mutations that escaped curation filters.

To investigate whether these variant classes experience differential selective pressures, predicted deleteriousness was quantified using the Combined Annotation Dependent Depletion (CADD) PHRED score, a composite metric integrating conservation, regulatory, and functional features into a single measure of pathogenic potential. CADD score distributions revealed a clear stratification across variant classes (Figure 2B). Homozygous- and heterozygous-like variants were strongly depleted for high CADD scores (≥20), with only 0.05–0.15% of variants exceeding this threshold, consistent with purifying selection against deleterious alleles at clonal frequency. In contrast, low-frequency variants exhibited a higher fraction (∼0.3%) of high-impact mutations, and somatic variants displayed the strongest enrichment (0.75%), suggestive of relaxed selection and ongoing mutational input. These results collectively support an evolutionary model in which germline-equivalent variants represent selectively filtered genomic backgrounds, whereas low-frequency and somatic mutations contribute a reservoir of potentially damaging, recently acquired changes.

### Functional annotation and mutational impact

*De novo* variants were further stratified by functional annotation to evaluate how mutational impact varies across genomic contexts. The majority of variants resided in intergenic and intronic regions, whereas exonic and splicing variants were two orders of magnitude rarer, consistent with genome composition. Despite their rarity, these coding-associated variants were markedly enriched for high CADD scores. Across HG001–HG005, 3–5% of exonic and nearly one-third of splicing variants exceeded the CADD PHRED threshold of 20, indicating substantial functional potential. Variants in untranslated regions showed intermediate CADD distributions (≤0.7% > 20), suggesting occasional effects on regulatory stability or transcript processing. In contrast, intergenic and intronic variants were overwhelmingly neutral (high-CADD < 0.1%). These patterns underscore that while most *de novo* variants occur in noncoding contexts, the subset within exonic and splice-related regions is disproportionately enriched for potentially impactful mutations.

### Potential biological and clinical relevance

Building upon the enrichment of high-CADD variants in coding regions, we next resolved the functional consequences of *de novo* variants in exons. Across HG001, HG002, and HG005, synonymous and missense substitutions comprise the majority of mutations and appear balanced, with only less than 1% mutations with strong impact (start-loss, stop-gain, or stop-loss, Figure 3A). High-impact variants are dispersed in individual genes rather than recurrent, consistent with purifying selection and the absence of in-vitro–driven clonal adaptation (Figure 3B). Concordantly, genes harboring predicted high-impact changes show minimal expression in blood cell lineages, indicating neutral-to-mild fitness effects in the GIAB lymphoblastoid culture and suggesting that the observed landscape reflects baseline somatic variation rather than culture selection.

**Figure 3.**
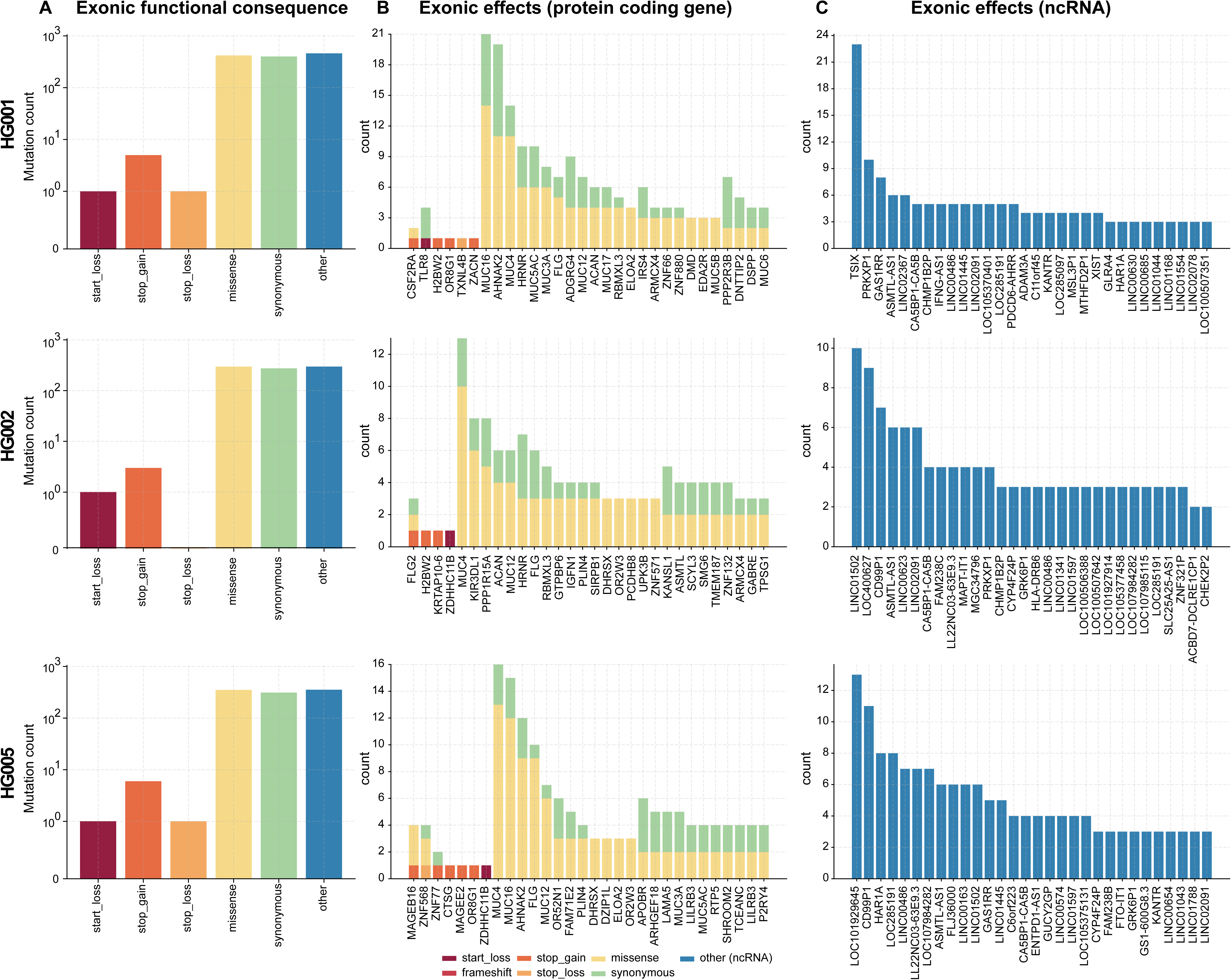
Functional annotation of de novo mutations. Rows: GIAB individuals (HG001, HG002, HG005). Columns: A) Exonic consequences (bar): counts by effect on protein coding. B) Gene attribution (bar): distribution of exonic effects across protein-coding genes. C) ncRNA attribution (bar): distribution of exonic effects across non-coding RNA genes.

Beyond coding sequence, we also observe ∼200–300 variants in UTRs or ncRNA genes (Figure 3C). Because such non-coding exonic elements can modulate transcript abundance, splicing, or RNA-mediated regulation under weaker protein-level constraint, they offer practical loci for rare-clone tracking and regulatory surveillance as sequencing scale increases.

Clinical relevance was assessed with OpenCRAVAT across ClinVar, ACMG, and OMIM resources. We identified 126 ClinVar-annotated variants in HG001, 82 in HG002, and 101 in HG005; none were classified as pathogenic, most were benign, and the remainder were variants of uncertain significance, with no ACMG pathogenic/likely pathogenic designations (Table S4-6). OMIM gene-level associations were observed for 24 genes in HG001, 16 in HG002, and 13 in HG005. Together with coverage of clinically relevant hotspots, these results indicate that the assay captures clinically interpretable variation while maintaining specificity, positioning it for early detection and prioritization of rare disease-associated somatic mutations.

### Somatic mutation burden and signatures in GIAB lymphoblastoid lines

Calibrating per-cell burden to each sample’s heterozygous germline detection rate, we estimate 694.3 (HG001), 620.3 (HG002), and 1050.7 (HG005) somatic mutations per cell (Figure 4A, Table S3). Although all three are immortalized lymphoblastoid cell lines (LCLs), HG005 shows a markedly higher burden. Base-substitution spectra point to a relative excess of C>T and T>G in HG005 (Figure 4B, Table S7). Decomposition of trinucleotide contexts against COSMIC reference signatures (Figure 4C, Table S8) indicates that SBS5 (clock-like) dominates all samples and is ∼200 mutations higher in HG005; SBS1—another replication/age-linked process—contributes 61.8 mutations per cell in HG005 versus 19 in HG002 and was not cleanly separable in HG001. Consistent with LCL culture, culture-associated signatures vary subtly: the total contribution of SBS18 and SBS36 is comparable in all three libraries.

**Figure 4.**
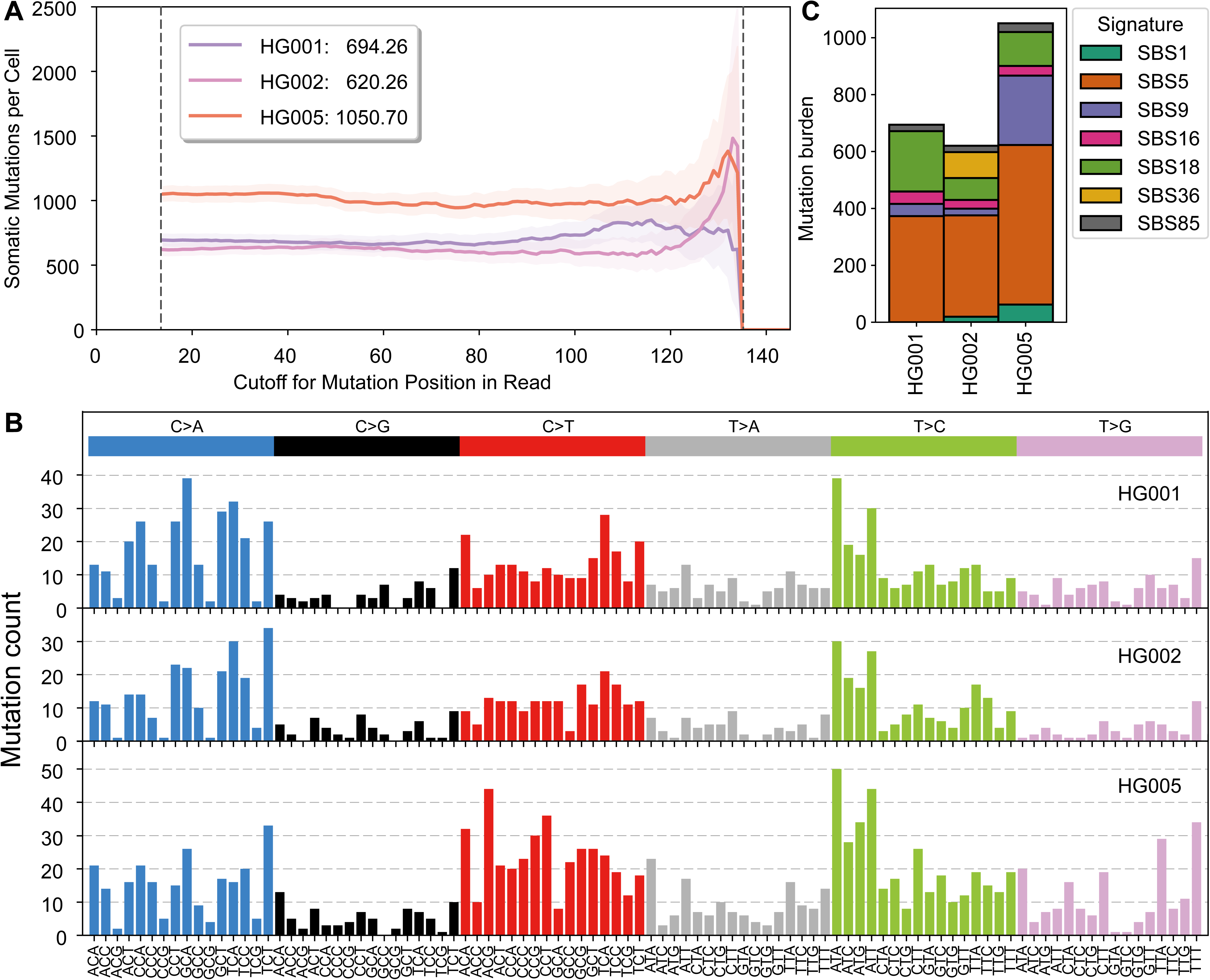
Somatic mutation burden and spectra across GIAB lymphoblastoid lines. (A) Per-cell somatic mutation burden across read-position cutoffs; all three samples plateau, indicating minimal position-specific bias. (B) Trinucleotide substitution spectra for HG001, HG002, and HG005. (C) Mutational-signature exposures normalized to somatic mutation burden. Cosine similarities between reconstructed and observed spectra are 0.943 (HG001), 0.951 (HG002), and 0.967 (HG005).

We also detect SBS9 in every sample—contributing 55 (HG001), 35 (HG002), and 260.7 (HG005) mutations per cell—implicating polymerase η–mediated off-target activity. Prior work (Machado et al.^17^, attributes SBS9 to polymerase η acting under elevated replicative stress and estimates that SBS9 events can outnumber on-target SHM mutations by ∼18-fold in normal memory B cells. This enrichment provides a parsimonious explanation for the near-absence of canonical SHM signatures (SBS84/85) in our LCLs: on-target activity is likely below detection even when offtarget SBS9 is measurable. Taken together, these patterns indicate that HG005’s increased burden reflects both stronger clock-like processes (SBS1 and SBS5) and elevated polymerase-η–driven mutagenesis, demonstrating that NanoSeq-MBN workflow enables accurate rare somatic mutation profiling, distinguishing mutagenesis contributed by various source.

## DISCUSSION

Low-frequency genetic variation is increasingly recognized as a pervasive layer of human diversity with direct relevance to disease biology ^18–20^. Somatic mosaicism— particularly variants with allele fractions well below conventional detection thresholds—contributes to neurodevelopmental phenotypes^21^, age-associated decline ^22^, and cancer risk ^23^, yet has remained underrepresented in benchmark resources. Here we optimized and demonstrate that NanoSeq-MBN with T4PNK workflow enables high-fidelity, near-genome-wide discovery of low-frequency and somatic variants and can operationally extend the GIAB resource from germline calibration to rare-variant and somatic benchmarking.

Using GIAB gold-standard samples (HG001, HG002, HG005), we built the three-sample mixture (HG005:HG001:HG002=60:2:1) and benchmarked multiple library strategies. NanoSeq-MBN delivered near–whole-genome coverage with minimal bias, and a low-mutation-burden cord-blood control confirmed low error rates. Single-sample surveys then revealed three classes of previously unreported variation: approximately 100,000 missed germline variants, around 3,000 low-frequency calls with attenuated bulk allele fractions, and 1,000 to 1,500 bona fide somatic mutations per genome (corresponding to 694 somatic mutations per cell in HG001, 620 in HG002, and 1,051 in HG005). Cross-validation against GIAB bulk datasets supports biological origin rather than culture or technical artifacts, indicating that duplex-level fidelity can uncover signals previously buried by sampling noise or context biases. ^14,16^.

Stratifying variants by allele fraction reveals selective and functional patterns across the genome. Germline-like variants show depletion of high-impact mutations, consistent with purifying selection, whereas low-frequency and somatic mutations exhibit relaxed constraint and evidence of recent clonal expansion. Functional annotation refines interpretation: although most events are noncoding, exonic and splice-proximal variants are disproportionately enriched for predicted deleteriousness. Clinical annotation further shows that NanoSeq-MBN faithfully recovers known benign and uncertain variants in lymphoblastoid lines without producing spurious pathogenic calls, confirming its high precision. Together, these findings underscore the research and preclinical value of NanoSeq-MBN’s genome-wide fidelity and motivate focused adaptations, such as targeted NanoSeq for coding regions. Beyond technical performance, integrating ultra-accurate somatic profiling into clinical genomics promises to refine genetic testing and counseling, illuminate variant mosaicism, and define the true baseline of somatic mosaicism—an essential step toward distinguishing disease-driving mutations from the neutral background of human variation ^24–26^.

Building on this foundation, NanoSeq-MBN has been implemented to support the Somatic Mosaicism across Human Tissues (SMaHT) network ^27–29^., enabling systematic discovery of ultra-rare variants across diverse tissues and paving the way for precision maps of human somatic diversity.

### Limitations and future directions

While duplex sequencing methods such as NanoSeq achieve exceptional accuracy for detecting ultra-rare single-nucleotide variants, their current design is not optimized for identifying small insertions and deletions, large structural variants, or copy-number alterations (shallow depth). Ongoing development aims to extend NanoSeq-MBN toward exome-enriched and hybrid-capture formats, enabling comprehensive assessment of coding regions and facilitating systematic discovery of functional or marker genes.

## Resource availability

## DATA AVAILABILITY

The raw sequencing data has been deposited in NCBI Sequence Read Archive (SRA) with BioProject accession code PRJNA1346471. GIAB Illumia HiSeq data was downloaded from the official GIAB FTP repository at https://ftptrace.ncbi.nlm.nih.gov/ReferenceSamples/giab/release/.

## CODE AVAILABILITY

The duplex-sequencing data were processed using the previously described analysis pipeline available at https://github.com/zonglab/CompDuplex. All custom scripts used for benchmarking, quality assessment, and comparative analyses are publicly available at https://github.com/Yang-Zhang-717/NanoSeq-MBN-Hapmap.

## ACKNOWLEDEMENTS

This work was supported by the National Institutes of Health (Grant# UM1DA058229 (R.A.G, H.D, and C.Z), and UG3NS132132 (C.Z)). No additional external funding was received for this study. The authors are grateful to the production teams at the Human Genome Sequencing Center for data generation.

## AUTHOR CONTRIBUTIONS

Conceptualization: H.D. Data Generation: H.C, C.M.G, K.K, S.V.B, Data Analysis: Y.Z. and M.N.; Writing: H.D, C.Z, Y.Z, H.C, M.N, C.M.G, K.K, Review & Editing: Y.Z, C.M.G, R.A.G, D.M.M, C.Z and H.D.

## DECLARATION OF INTRESTS

The authors declare no conflict of interest.

## STAR ★ METHODS

Detailed methods are provided in the online version of this paper and include the following:

## KEY RESOURCE TABLE

## METHOD DETAILS

### a. Samples

DNA for the three benchmarking samples, HG001 (NA12878, Female), HG002 (NA24385, Male), and HG005 (NA24631, Male) were ordered from Coriell Institute for Medical Research and human cord blood sample was ordered from STEM CELL TECHNOLOGIES (Catalog # 70007.2). DNA was initially quantified using qubit. Based on this QC, samples were next diluted to 10ng/ul, and another round of qubit was performed on these aliquots for added precision. “Along with Qubit QC, qPCR was performed using RPL30 primers (5’-GCCCGTTCAGTCTCTTCGATT forward, 5’CAAGGCAAAGCGAAATTGGT reverse, ^30^) on individual samples to ensure consistency in the amount of DNA used to prepare the spike-in mixtures. To set up qPCR, reaction mixture (10 μL) included 5 μL of KAPA SYBR FAST master mix (KAPA library quantification kit (KK4835)), 0.25 μL of forward and reverse primer (20 μM) and 50ng of DNA template. Purified standards and no-template controls were also included in all qPCR runs. Thermal cycling parameters were as follows: hold at 95°C for 10 min to achieve initial denaturation, followed by 40 cycles of: 10 s hold at 95°C to denature, ramp-down to 60°C for primer annealing and extension occurring through a 35s rampup to 95°C, all the reactions were set up in triplicates.

For three-sample mixes, HG001 and HG002 were first combined in a 2:1 ratio to a total of 1 µg and thoroughly mixed by slow pipetting. From this 2:1 (HG001:HG002) mixture, an additional mixing was performed with HG005 in a 20:1 ratio (HG005: twosample mix), yielding a final 3 sample mix spike-in ratio of 60:2:1 (HG005: HG001: HG002).

### b. NanoSeq Sequencing data generation

#### NanoSeq-RE Library

To prepare the NanoSeq-RE libraries, three 200 ng DNA aliquots of the mix were fragmented in 25 μl 1X CutSmart buffer. DNA from one aliquot was fragmented using 5 units HpyCh4V enzyme, the second aliquot in 5 units of AluI enzyme, and the third using 5 units of Hpy166II enzyme. Fragmented DNA was purified with 2X AMPure XP beads and eluted in 40 μl nuclease-free water. Three enzymatic NanoSeq libraries were then prepared from different combinations of fragmented DNA: (1) 10 μl HpyCh4V-fragmented DNA, (2) 5 μl each of HpyCh4V- and AluI-fragmented DNA, and (3) 3.3 μl each of HpyCh4V-, AluI-, and Hpy166II-fragmented DNA. The DNA inputs for these libraries were approximately 50 ng. Libraries were prepared in duplicate for each combination of restriction enzyme group. Fragmented DNA samples were subjected to A-tailing and adapter ligation following the protocol described by Abascal *et al* ^7^. Adapter-ligated libraries were purified twice with 1X AMPure XP beads and eluted in 10 μl Tris buffer, then diluted 10-fold and quantified by qPCR.

#### NanoSeq-MBN library

Fifty nanograms DNA aliquots of the three-sample mixture and cord blood samples were used to prepare Mung Bean Nuclease NanoSeq (MBN) libraries. Duplicate libraries were prepared for each sample as described by Chao et al. ^31^, and two additional libraries were prepared using the same protocol but without the T4PNK treatment. Briefly, DNA were sheared, purified then treated with 0.5-unit Mung Bean nuclease. Samples were purified with 2.0X AMPure XP beads and eluted into 11 μl NFW. For libraries with T4PNK treatment, 5 μl of master mix of 1.5 μl NEBuffer™ 4 (NEB, B7004S), 1 μl NFW, 1.5 μl 10 mM ATP and 1 μl T4PNK was added to 10 μl MBN treated DNA and then were incubated at 37°C for 30 min on the thermocycler.

For 2 libraries without T4PNK treatment, 5 μl of master mix of 1.5 μl NEBuffer™ 4 (NEB, B7004S), 3.5 μl NFW was added to sample and held on ice during the T4PNK incubation. A-Tailing and adapter ligation were performed as described in the reference protocol. Adapter-ligated libraries were purified twice with 1X× AMPure XP beads and eluted in 25 μl Tris buffer, then diluted 20-fold and quantified by qPCR.

Six NanoSeq-MBN libraries were generated in duplicate using 50 ng DNA from NA12878, NA24385, and NA24631, following the referenced protocol without modification.

### Post-Adapter Ligation Library qPCR Quantification

The library qPCR master mix was prepared by adding primer premix provided with the kit and 20 μl of 100 μM Nano qPCR primer 1 & 2 (HPLC, 5’ACACTCTTTCCCTACACGAC-3’, HPLC, 5’-GTGACTGGAGTTCAGACGTG-3’) mix to KAPA SYBR FAST master mix. Library qPCR was performed in a 10 μl reaction containing 6 μl of qPCR master mix, 2 μl sample (20x dilution) and 2 μl NFW. Reactions were set as triplicates in 384 well plates and were run on QuantStudio™ 6 Flex Real-Time PCR System (Applied Biosystems™). The library concentration (nM) is determined by taking the average library size as 573 bp. Based on qPCR result, 0.1 fmol library was made into 23.5 μl with NFW for final library amplification.

### Library Amplification

NanoSeq libraries were amplified in a 50 μl PCR reaction containing post-adapter ligated library, 1.5 μl xGen™ UDI 10nt primer mix (IDT, 10008052) and 25 μl of NEBNext Ultra II Q5 Master Mix. The PCR reactions were performed in the thermocycler program: step 1, 98°C 30s; step 2, 98°C 10s; step 3, 65°C 75s; step 4, return to step 2 for *N* times; step 5, 65°C for 5 min; step 6, hold at 4°C. For enzymatic NanoSeq libraries, 0.3 fmol libraries were amplified for 14 PCR cycles. For MBN NanoSeq libraries, 0.1 fmol libraries were amplified for 16 PCR cycles. Amplified libraries were purified by adding 30 μl NFW and 72 μl AMPure XP beads and eluted in 20 μl Tris buffer. Purified libraries were QC’ed on DNA 7500 kit (Agilent, 5067-1506). Libraries were normalized to 10 nM and quantified using qPCR KAPA library quantification kit. Libraries were pooled and sequenced on Illumina NovaSeq sequencing platforms for 30x coverage.

### Whole genome sequencing

PCR-free libraries were prepared to generate short-read WGS data for HG001 (NA12878, Female), HG002 (NA24385, Male), HG005 (NA24631, Male) and human cord blood samples using the KAPA Hyper Prep Kit as previously described ^32,33^. DNA (0.75 μg) was sheared into fragments of approximately 450-600 bp in a Covaris E220 system (Covaris, Inc. Woburn, MA), followed by double SPRI bead cleanup with different ratios of AMPure XP beads to select a narrow band of sheared DNA for library preparation. Using KAPA Hyper kit (KAPA Biosystems Inc., #KK8505), DNA endrepair and 3’-adenylation are performed in the same reaction followed by ligation of the barcoded adaptors to create PCR-Free libraries. A set of 96 8-bp barcoded adapters (Illumina, TruSeq UD Indexes v2, # 20040870) were used for sample barcoding. Post ligation products are cleaned twice with AMPure XP to get rid of leftover adapters/adapter dimers that may otherwise interfere with sequencing. The final library sizes and quantities were estimated using the Fragment Analyzer (Agilent Advanced Analytical Technologies, Inc) electrophoresis system and Quant Studio 6 Flex Real-Time PCR System (Applied Biosystem) respectively. Samples were pooled at equimolar ratios and sequenced on the NovaSeq X platform using a 25B flow cell (2×150 bp) for short-read sequencing to achieve 30x coverage.

### NanoSeq sequencing data analysis

All primary processing was executed on a Slurm HPC cluster under a unified analysis pipeline for all NanoSeq libraries. Raw FASTQ files were adapter-trimmed with cutadapt v5.0 (minimum read length 50; -a/-A CTGTCTCTTATACACATCT) followed by optional tag extraction (for UMI/inline tags) using a customized Python script.

Reads were aligned to GRCh38 (GCA_000001405.15, no-alt analysis set) using BWAMEM v0.7.13-r1126 (-C -K 1000000000 with read-group tags) and converted to BAM with samtools. Optical/sequence duplicate handling and coordinate sorting were performed as in the workflow; when applied, duplicates were marked/removed with Picard Mark Duplicates v3.4.0 (queryname sort; REMOVE_SEQUENCING_DUPLICATES=true), and quality filtering used samtools view -q 50 -f 3 prior to final sorting and indexing.

Per-chromosome variant discovery first enumerated candidate sites from aligned reads using samtools v1.12 mpileup (excluding QCFAIL) piped to bcftools v1.22 with a site filter requiring sufficient alternate evidence ((I16[2] +I16[3]) > 3). Multiallelic normalization and left alignment were performed with bcftools (-m-any -f GRCh38), and symbolic alleles were removed. Ambiguous non-ACGT records were filtered with a customized Python script. Variants overlapping the Abascal genome mask were excluded via bedtools v2.31.1 intersect -v against the consolidated blacklist BED.

Double-strand consensus reconstruction used customized Python scripts to collapse read-pairs into duplex molecules and to apply an aMsN-style consensus-aware caller for SNVs, parameterized by strand/read support and read-length thresholds. These scripts take the candidate site lists and the per-chromosome BAMs as input and output high-confidence consensus calls for both SNV and indel classes.

To separate true germline from de novo/somatic events, we demultiplexed detected variants against matched bulk BAMs and retained only variants absent from bulk. Remaining calls were annotated with SnpSift (from the snpEff suite) against dbSNP resources (Broad dbSNP138 20180418) and any record carrying an rsID tag was removed to eliminate missed germline contamination.

Variant records from duplex consensus calling were parsed from tabular callsets with standardized quality filters. We retained sites with properly bounded paired-inter-read (PIR) values, sufficient fragment length, robust alignment score separation (AS–XS), and adequate Phred-30 base fraction: PIR between 8 and 136, insert size ≥100 bp, AS–XS ≥50, and *Q*_30_/(*Q*_30_ + *Q*_low_) ≥ 0.5.INFO fields were parsed to extract auxiliary metrics; an optional “bulk” column encodes “DENOVO/BULKCOV” for cross-checking in bulk alignments (below).

### Analysis of the original GIAB samples sequence data to confirm the variants

To orthogonally validate duplex-seq discoveries and quantify background variation, we reanalyzed the original bulk HiSeq data for HG001, HG002, and HG005. We regenerated read pairs from the archived BAMs and realigned to GRCh38 (GCA_000001405.15, no-alt analysis set).

Adapter trimming used cutadapt v5.0 (-m 50 -a/-A CTGTCTCTTATACACATCT), optionally preceded by UMI/inline-tag extraction with a customized Python script when tags were present. Trimmed reads were aligned to GRCh38 (no-alt) with BWA-MEM v0.7.13-r1126 (-C -K 1000000000) and read-group tags. BAM conversion and initial processing used samtools; duplicates were marked/removed with Picard MarkDuplicates v3.4.0 (queryname sort; REMOVE_SEQUENCING_DUPLICATES=true). We applied a stringent mapping-quality/read-pair filter prior to final coordinate sorting and indexing (samtools view -q 50 -f 3).

Post-alignment, we split the deduplicated BAM by chromosome (samtools), indexed each chromosome, and applied GATK v4.5.0.0 BaseRecalibrator/ApplyBQSR using dbSNP (Broad dbSNP138 20180418) as known sites, followed by Haplotype Caller to produce VCFs per chromosome.

VCFs were decompressed (pigz) and normalized with bcftools v1.22 norm (-m any, left-align against the same GRCh38 reference). We then applied hard filters: SNPs required QUAL>30 and ≥5 alternate reads (FORMAT/AD) and were written to per-chromosome SNV VCFs.

Per-chromosome SNV calls were aggregated and re-headed to produce a wholegenome SNV VCF. To remove problematic loci, we excluded sites overlapping either the GIAB composite genome mask or the NanoSeq mask using bedtools v2.31.1 intersect -v.

For additional sequence-context QC, we extracted ±10 bp around each site with seqtk v1.4-r122 subseq and passed the VCF plus context to a custom Python local-sequence filter to generate “clean” outputs. Cleaned calls were separated by genotype into heterozygous and homozygous sets and reassembled with the preserved VCF header.

### Genotype-aware classes and expected allele fractions for three-sample mixture

We constructed a genotype-aware expectation for each variant site using donor-specific genotypes and the known mass fractions of each donor in the mixture. For each locus *l*, let *d*_*i*_(*l*) ∈ {0,1,2} denote the alternate-allele dosage for donor *i*, and *w*_*i*_ the donor’s DNA mass fraction (∑ *w*_*i*_ = 1). The expected VAF in the mixture is

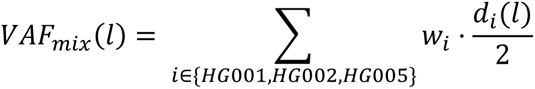

Using per-donor genotypes, loci were grouped into genotype-aware classes (e.g., “heterozygous-only”, “homozygous-only”, or composite classes spanning multiple donors) and, for each class, we aggregated (i) the number of loci, (ii) the class-wise expected VAF *VAF*_*mix*_(*l*), and (iii) the heterozygous- and homozygous-specific expected contributions.

### Expected duplex-molecule counts and origin-constrained calibration

For each library, we enumerated observed duplex molecules that supported the alternate allele at truth loci and summed these counts by genotype-aware class.

Expected counts at nominal “1× molecular depth” equal the class-wi by regressing observed*V*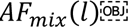 on expected counts 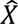 by regressing observed counts *y*_*c*_ on expected counts *x*_*c*_ across classes with a no-intercept lease-squares fit:

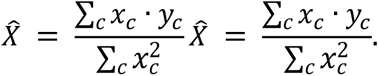

Uncertainty in 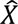 was quantified by bootstrap resampling across classes. The analysis function merges expected and observed class totals, computes 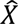 from dot-product ratio and bootstraps 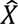.

### Per-locus molecule counts and zero-inflated negative binomial (ZINB) coverage model

To model variability in per-locus duplex-molecule counts, we fit a zero-inflated negative binomial (ZINB) distribution to the empirical count distribution within each class. The ZINB has a structural-zero probability *π*and, conditional on being nonstructural, follows a negative binomial with mean *μ*_NB_and dispersion *θ*. The negativebinomial mass for *k*molecules is

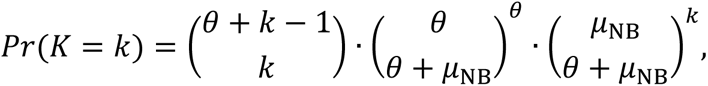

and Pr (*K* = 0) = *π* + (1 − *π*) Pr _NB_(0). Parameters are estimated by moment-matching: we enumerate a grid of *θ*values and, for each, select *μ*_NB_so that the model matches the observed mean; *π*is then chosen to match the observed zero fraction. We report 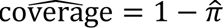 as the fraction of loci with at least one observed duplex molecule, with bootstrap confidence intervals obtained by resampling per-locus counts.

### Somatic mutation burden from duplex molecules

We define *mutation count by molecule* at locus *l*as the number of independent duplex molecules in which the consensus sequence supports the alternate allele. Somatic burden per library was inferred by normalizing the number of *somatic* duplex-supported events to the number of duplex-supported *germline heterozygous* events and then scaling by the total number of heterozygous alleles in the evaluated genome segment:

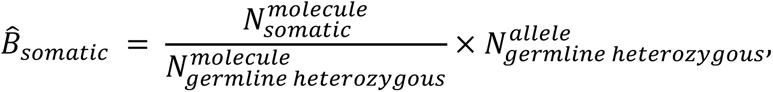

where 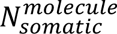 and 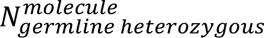 are the molecule-level counts observed in the duplex data for somatic and germline heterozygous mutations, and 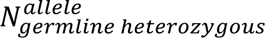 is the number of truth-set heterozygous alleles within the callable space. This estimator leverages germline heterozygous recovery as an internal calibration of effective molecular depth.

### Identification of de novo candidates and cross-check in GIAB bulk data

We enumerated de novo candidates by removing every event that matched a truthset key (chromosome, position, reference, alternate) and counting unique duplex-supported events among the remaining records; per-event molecule counts are tracked for downstream modeling. For records with a “bulk” field, the pipeline preserves “DENOVO/BULKCOV” summaries derived from the original BAMs to facilitate manual re-examination.

### VAF mixture modeling for class assignment

To model the distribution of VAFs across putative de novo and germline events, we employed a logit-normal finite mixture with an explicit point mass near zero. Specifically, after separating a “zero-spike” proportion *w*_0_ = Pr (VAF ≤ *ɛ*) with *ɛ* = 0.005, the remaining continuous density is represented as a non-negative leastsquares combination of three logit-normal components centered near ∼0.15, ∼0.50, and ∼0.98 for low-frequency, heterozygous, and homozygous modes, respectively. Grid-searched component standard deviations minimize mean-squared error to a binned density estimate. Posterior class probabilities are computed as *P*(*c* ∣ *v*) ∝ *w*_*c*_*f*_*c*_(*v*)and used for per-site class labels.

### Functional annotation and pathogenicity scoring

Functional consequences were assigned using ANNOVAR against human reference gene models; pathogenicity was summarized with Combined Annotation-Dependent Depletion (CADD) PHRED-scaled scores. Statistical summaries included nonparametric tests across genotype classes and enrichment of high-tail CADD scores within coding-like regions (exonic, splicing, untranslated regions), with optional visualization of distributional differences and high-tail proportions.

### Assess the potential biological and clinical relevance

Coding sequence variants from three reference genomes (HG001, HG002, and HG005) were analyzed using the Open Custom Ranked Analysis of Variants Toolkit ^34^(CRAVAT) to annotate functional effects and their clinical relevance. We assessed ClinVar (Hg38) annotations ^35,36^, as well as Online Mendelian Inheritance in Man (OMIM) gene associations. ClinVar entries were extracted to evaluate the number and classification of variants with prior clinical interpretation. OMIM associations reflect gene-level links to Mendelian disorders, not variant-specific disease correlations.

## SUPPLEMENTAL INFORMATION

## REFERENCES

1. Bizzotto, S. (2023). The human brain through the lens of somatic mosaicism. Frontiers in Neuroscience Volume 17-2023. 10.3389/fnins.2023.1172469.

2. Ganz, J., Luquette, L.J., Bizzotto, S., Miller, M.B., Zhou, Z., Bohrson, C.L., Jin, H., Tran, A.V., Viswanadham, V.V., McDonough, G., et al. (2024). Contrasting somatic mutation patterns in aging human neurons and oligodendrocytes. Cell 187, 1955-1970.e1923. 10.1016/j.cell.2024.02.025.

3. Ha, Y.-J., Kang, S., Kim, J., Kim, J., Jo, S.-Y., and Kim, S. (2023). Comprehensive benchmarking and guidelines of mosaic variant calling strategies. Nature Methods 20, 2058–2067. 10.1038/s41592-023-02043-2.

4. Pareja, F., Ptashkin, R.N., Brown, D.N., Derakhshan, F., Selenica, P., da Silva, E.M., Gazzo, A.M., Da Cruz Paula, A., Breen, K., Shen, R., et al. (2022). Cancer-Causative Mutations Occurring in Early Embryogenesis. Cancer Discovery 12, 949–957. 10.1158/2159-8290.Cd-21-1110.

5. Menon, V., and Brash, D.E. (2023). Next-generation sequencing methodologies to detect low-frequency mutations: “Catch me if you can”. Mutat Res Rev Mutat Res 792, 108471. 10.1016/j.mrrev.2023.108471.

6. Axelsson, J., LeBlanc, D., Shojaeisaadi, H., Meier, M.J., Fitzgerald, D.M., Nachmanson, D., Carlson, J., Golubeva, A., Higgins, J., Smith, T., et al. (2024). Frequency and spectrum of mutations in human sperm measured using duplex sequencing correlate with trio-based de novo mutation analyses. Scientific Reports 14, 23134. 10.1038/s41598-024-73587-2.

7. Abascal, F., Harvey, L.M.R., Mitchell, E., Lawson, A.R.J., Lensing, S.V., Ellis, P., Russell, A.J.C., Alcantara, R.E., Baez-Ortega, A., Wang, Y., et al. (2021). Somatic mutation landscapes at single-molecule resolution. Nature 593, 405–410. 10.1038/s41586-021-03477-4.

8. Schmitt, M.W., Kennedy, S.R., Salk, J.J., Fox, E.J., Hiatt, J.B., and Loeb, L.A. (2012). Detection of ultra-rare mutations by next-generation sequencing. Proc Natl Acad Sci U S A 109, 14508–14513. 10.1073/pnas.1208715109.

9. Kennedy, S.R., Schmitt, M.W., Fox, E.J., Kohrn, B.F., Salk, J.J., Ahn, E.H., Prindle, M.J., Kuong, K.J., Shen, J.C., Risques, R.A., and Loeb, L.A. (2014). Detecting ultralow-frequency mutations by Duplex Sequencing. Nat Protoc 9, 2586–2606. 10.1038/nprot.2014.170.

10. Cheng, A.P., Rusinek, I., Sossin, A., Widman, A.J., Meiri, E., Krieger, G., Hirschberg, O., Tov, D.S., Gilad, S., Jaimovich, A., et al. (2025). Paired plus-minus sequencing is an ultra-high throughput and accurate method for dual strand sequencing of DNA molecules. bioRxiv. 10.1101/2025.08.11.669689.

11. Bae, J.H., Liu, R., Roberts, E., Nguyen, E., Tabrizi, S., Rhoades, J., Blewett, T., Xiong, K., Gydush, G., Shea, D., et al. (2023). Single duplex DNA sequencing with CODEC detects mutations with high sensitivity. Nat Genet 55, 871–879. 10.1038/s41588-023-01376-0.

12. You, X., Cao, Y., Suzuki, T., Shao, J., Zhu, B., Masumura, K., Xi, J., Liu, W., Zhang, X., and Luan, Y. (2023). Genome-wide direct quantification of in vivo mutagenesis using high-accuracy paired-end and complementary consensus sequencing. Nucleic Acids Research 51, e109–e109. 10.1093/nar/gkad909.

13. Troll, C.J., Kapp, J., Rao, V., Harkins, K.M., Cole, C., Naughton, C., Morgan, J.M., Shapiro, B., and Green, R.E. (2019). A ligation-based single-stranded library preparation method to analyze cell-free DNA and synthetic oligos. BMC Genomics 20, 1023. 10.1186/s12864-019-6355-0.

14. Zook, J.M., Chapman, B., Wang, J., Mittelman, D., Hofmann, O., Hide, W., and Salit, M. (2014). Integrating human sequence data sets provides a resource of benchmark SNP and indel genotype calls. Nat Biotechnol 32, 246–251. 10.1038/nbt.2835.

15. Zook, J.M., Catoe, D., McDaniel, J., Vang, L., Spies, N., Sidow, A., Weng, Z., Liu, Y., Mason, C.E., Alexander, N., et al. (2016). Extensive sequencing of seven human genomes to characterize benchmark reference materials. Sci Data 3, 160025. 10.1038/sdata.2016.25.

16. Zook, J.M., McDaniel, J., Olson, N.D., Wagner, J., Parikh, H., Heaton, H., Irvine, S.A., Trigg, L., Truty, R., McLean, C.Y., et al. (2019). An open resource for accurately benchmarking small variant and reference calls. Nat Biotechnol 37, 561–566. 10.1038/s41587-019-0074-6.

17. Machado, H.E., Mitchell, E., Øbro, N.F., Kübler, K., Davies, M., Leongamornlert, D., Cull, A., Maura, F., Sanders, M.A., Cagan, A.T.J., et al. (2022). Diverse mutational landscapes in human lymphocytes. Nature 608, 724–732. 10.1038/s41586-02205072-7.

18. Wright, C.F., Prigmore, E., Rajan, D., Handsaker, J., McRae, J., Kaplanis, J., Fitzgerald, T.W., FitzPatrick, D.R., Firth, H.V., and Hurles, M.E. (2019). Clinically-relevant postzygotic mosaicism in parents and children with developmental disorders in trio exome sequencing data. Nat Commun 10, 2985. 10.1038/s41467-019-11059-2.

19. Miller, K.E., Rivaldi, A.C., Shinagawa, N., Sran, S., Navarro, J.B., Westfall, J.J., Miller, A.R., Roberts, R.D., Akkari, Y., Supinger, R., et al. (2023). Post-zygotic rescue of meiotic errors causes brain mosaicism and focal epilepsy. Nat Genet 55, 1920-1928. 10.1038/s41588-023-01547-z.

20. Hsieh, A., Morton, S.U., Willcox, J.A.L., Gorham, J.M., Tai, A.C., Qi, H., DePalma, S., McKean, D., Griffin, E., Manheimer, K.B., et al. (2020). EM-mosaic detects mosaic point mutations that contribute to congenital heart disease. Genome Med 12, 42. 10.1186/s13073-020-00738-1.

21. D’Gama, A.M., and Walsh, C.A. (2018). Somatic mosaicism and neurodevelopmental disease. Nat Neurosci 21, 1504–1514. 10.1038/s41593-018-0257-3.

22. Martincorena, I., Fowler, J.C., Wabik, A., Lawson, A.R.J., Abascal, F., Hall, M.W.J., Cagan, A., Murai, K., Mahbubani, K., Stratton, M.R., et al. (2018). Somatic mutant clones colonize the human esophagus with age. Science 362, 911–917. 10.1126/science.aau3879.

23. Yizhak, K., Aguet, F., Kim, J., Hess, J.M., Kubler, K., Grimsby, J., Frazer, R., Zhang, H., Haradhvala, N.J., Rosebrock, D., et al. (2019). RNA sequence analysis reveals macroscopic somatic clonal expansion across normal tissues. Science 364. 10.1126/science.aaw0726.

24. Coban-Akdemir, Z.H., Charng, W.L., Azamian, M., Paine, I.S., Punetha, J., Grochowski, C.M., Gambin, T., Valdes, S.O., Cannon, B., Zapata, G., et al. (2020). Wolff-Parkinson-White syndrome: De novo variants and evidence for mutational burden in genes associated with atrial fibrillation. Am J Med Genet A 182, 1387-1399. 10.1002/ajmg.a.61571.

25. Gambin, T., Liu, Q., Karolak, J.A., Grochowski, C.M., Xie, N.G., Wu, L.R., Yan, Y.H., Cao, Y., Coban Akdemir, Z.H., Wilson, T.A., et al. (2020). Low-level parental somatic mosaic SNVs in exomes from a large cohort of trios with diverse suspected Mendelian conditions. Genet Med 22, 1768–1776. 10.1038/s41436-020-0897-z.

26. Pehlivan, D., Bayram, Y., Gunes, N., Coban Akdemir, Z., Shukla, A., Bierhals, T., Tabakci, B., Sahin, Y., Gezdirici, A., Fatih, J.M., et al. (2019). The Genomics of Arthrogryposis, a Complex Trait: Candidate Genes and Further Evidence for Oligogenic Inheritance. Am J Hum Genet 105, 132–150. 10.1016/j.ajhg.2019.05.015.

27. Coorens, T.H.H., Oh, J.W., Choi, Y.A., Lim, N.S., Zhao, B., Voshall, A., Abyzov, A., Antonacci-Fulton, L., Aparicio, S., Ardlie, K.G., et al. (2025). The Somatic Mosaicism across Human Tissues Network. Nature 643, 47–59. 10.1038/s41586-025-09096-7.

28. Zhang, Y., group, S.d.f., and Coorens, T. (2025). Benchmarking of duplex sequencing approaches to uncover somatic mutation landscapes.

29. Network, T.S.M.a.H.T. (2025). Comprehensive benchmarking of somatic mutation detection by the SMaHT Network.

30. D’Amico, L., Ajami, N.J., Adachi, J.A., Gascoyne, P.R., and Petrosino, J.F. (2017). Isolation and concentration of bacteria from blood using microfluidic membraneless dialysis and dielectrophoresis. Lab Chip 17, 1340–1348. 10.1039/c6lc01277a.

31. Hsu, C., Kottapalli, K., Jiang, Q., Niu, M., Zong, C.C., and Doddapaneni, H. (2025). Optimized Mung Bean Nuclease NanoSeq Library Preparation Protocol for Whole Genome Sequencing.

32. Hill, E.J., Robak, L.A., Al-Ouran, R., Deger, J., Fong, J.C., Vandeventer, P.J., Schulman, E., Rao, S., Saade, H., Savitt, J.M., et al. (2022). Genome Sequencing in the Parkinson Disease Clinic. Neurol Genet 8, e200002. 10.1212/NXG.0000000000200002.

33. McDaniel, J.H., Patel, V., Olson, N.D., He, H.J., He, Z., Cole, K.D., Gooden, A.A., Schmitt, A., Sikkink, K., Sedlazeck, F.J., et al. (2025). Development and extensive sequencing of a broadly-consented Genome in a Bottle matched tumor-normal pair. Sci Data 12, 1195. 10.1038/s41597-025-05438-2.

34. Pagel, K.A., Kim, R., Moad, K., Busby, B., Zheng, L., Tokheim, C., Ryan, M., and Karchin, R. (2020). Integrated Informatics Analysis of Cancer-Related Variants. JCO Clin Cancer Inform 4, 310–317. 10.1200/CCI.19.00132.

35. Rehm, H.L., Berg, J.S., Brooks, L.D., Bustamante, C.D., Evans, J.P., Landrum, M.J., Ledbetter, D.H., Maglott, D.R., Martin, C.L., Nussbaum, R.L., et al. (2015). ClinGen--the Clinical Genome Resource. N Engl J Med 372, 2235–2242. 10.1056/NEJMsr1406261.

36. Wilcox, E.H., Webb, R.F., Tshering, K.C., Hughes, M.Y., Cave, H., DiStefano, M.T., Dziadzio, H., Garber, K., Gelb, B.D., Gripp, K.W., et al. (2025). Updated ACMG/AMP specifications for variant interpretation and gene curations from the ClinGen RASopathy expert panels. Genet Med Open 3, 103430. 10.1016/j.gimo.2025.103430.

