## Supplementary Notes for "Expanding the Genome in a Bottle Truth Set: Detection and Validation of Novel Low-frequency Variants Using High-accuracy NanoSeq"

**Supplementary Note 1. Library quality control metrics.**
Enzymatic sequencing libraries prepared from the three-sample mixture (HG005:HG001:HG002=60:2:1) exhibited high alignment rates (99.7–99.8%) with median insert sizes averaging approximately 250 bp. Mung Bean NanoSeq (MBN) libraries generated with and without T4 polynucleotide kinase (T4PNK) yielded highly comparable results, showing mean insert sizes of roughly 236 bp. When the MBN + T4PNK workflow was applied to individual Genome-in-a-Bottle reference samples (HG001, HG002, and HG005), alignment rates ranged from ~95% to ~99%, indicating robust and consistent library performance across diverse genomic backgrounds. Whole-genome sequencing (WGS) of these individual samples produced longer inserts, with mean fragment sizes averaging 414 bp.

**Supplementary Fig. 1. Extended UpSet plot summarizing genotype-aware mutation classification across library designs. (A–E)** (A–E) Stacked histograms show the per-class distribution of expected versus observed molecule detections across the five NanoSeq strategies. Each panel corresponds to one library design, with bars grouped by genotype category (heterozygous, homozygous, or *de novo*) and colored by genotype contribution (total mutations, white; detected heterozygous mutations, blue; detected homozygous mutations, red). The dot centered on each bar indicates the number of detected mutations per class. The x-axis enumerates the 26 genotype-defined mutation classes derived from the GIAB trio, plus *de novo* mutations, and the y-axis represents molecule counts. (F) Lower left bar plot reflects mutation counts in corresponding donors stratified by genotype. (G) The accompanying dot plot illustrates class combinations shared among donors, with dot color indicating the genotype status in each donor and dot size proportional to the total number of loci per combination. Across all library designs, observed detections closely match theoretical expectations, demonstrating chemistry-independent fidelity of duplex consensus calling and validating the use of genotype-aware calibration for downstream analyses.

**Supplementary Fig. 2. Genomic coverage and trinucleotide context profiles of five NanoSeq optimization designs. (A, B)** Histograms show per-locus *molecule coverage*—defined as the number of unique duplex molecules spanning each locus, in contrast to *read coverage*, which counts total sequencing reads regardless of molecular origin. **(A)** All unfiltered molecules; **(B)** high-fidelity molecules passing the a4s2 filter. Zero-inflated negative binomial (ZINB) fits align almost perfectly with empirical distributions, indicating that the model's over-dispersion parameter accurately captures locus-to-locus variability introduced during library preparation. $\pi$: fraction of technically unsequenceable genome; $\mu$: mean molecule depth; $\alpha$: over-dispersion relative to the Poisson expectation. **(C)** Trinucleotide-context enrichment relative to the GRCh38 genomic baseline. Bars represent fold-enrichment (y-axis) across all 96 trinucleotide contexts, with the legend reporting the standard deviation across contexts as a global measure of sequence bias. Bars with height below 1 indicate depletion of specific motifs. For example, NanoSeq-HpyCH4V exhibits strong depletion of GCA contexts corresponding to the HpyCH4V restriction-enzyme cut site, and pronounced GC enrichment likely reflecting PCR-amplification bias. While both MBN libraries present flat profiles, consistent with near Poisson coverage for even distribution.

**Table S1. NanoSeq libraries**
**Table S2. Duplex analysis metrics for three-sample mix**
**Table S3. Duplex analysis metrics for single samples**
**Table S4. Clinically relevant mutations HG001**
**Table S5. Clinically relevant mutations HG002**
**Table S6. Clinically relevant mutations HG005**
**Table S7. Somatic mutation trinucleotide profiles**
**Table S8. Somatic mutation signature exposures**
