## Supplementary figures and images for "Expanding the Genome in a Bottle Truth Set: Detection and Validation of Novel Low-frequency Variants Using High-accuracy NanoSeq"

### Supplementary Figure 1

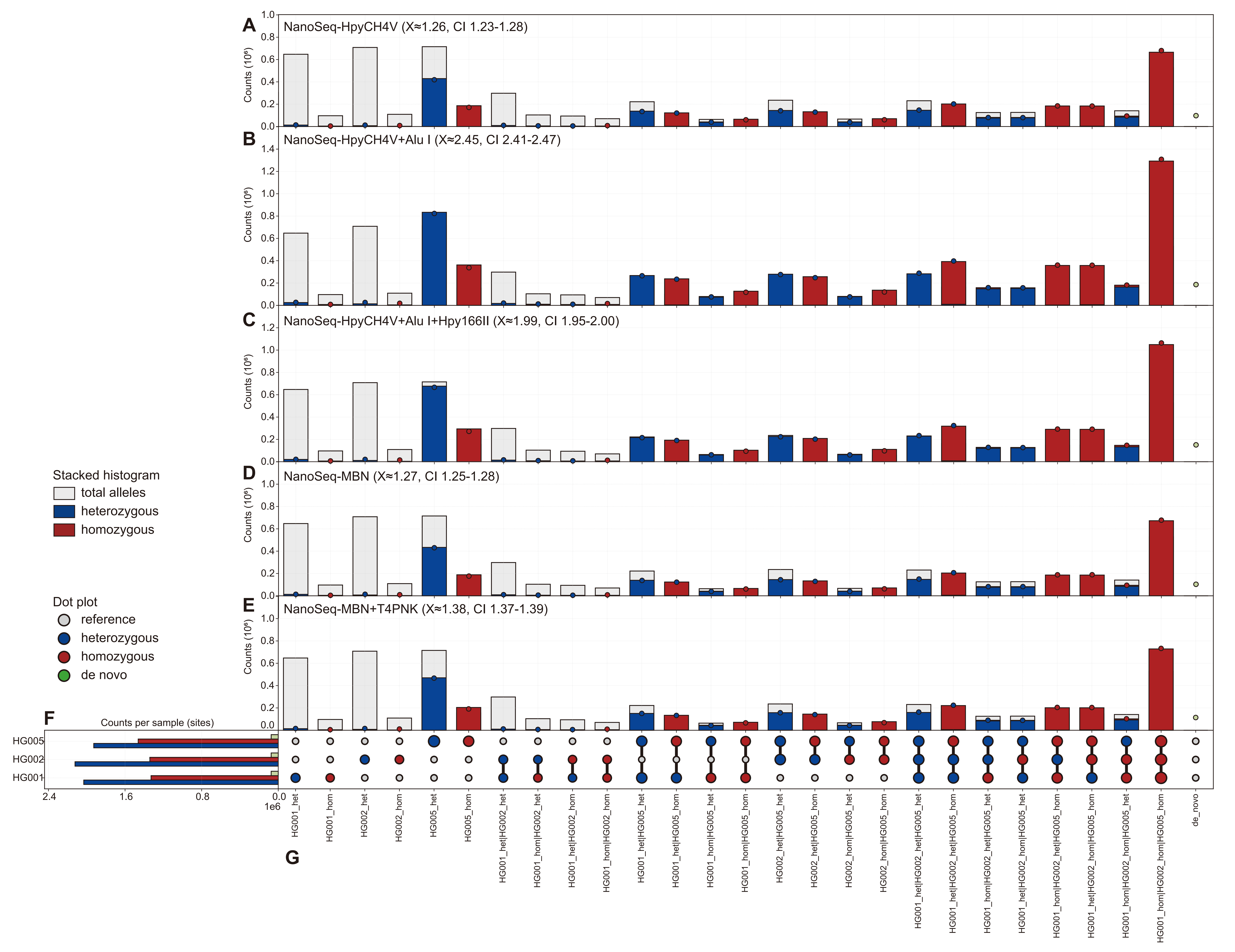

### Supplementary Figure 2

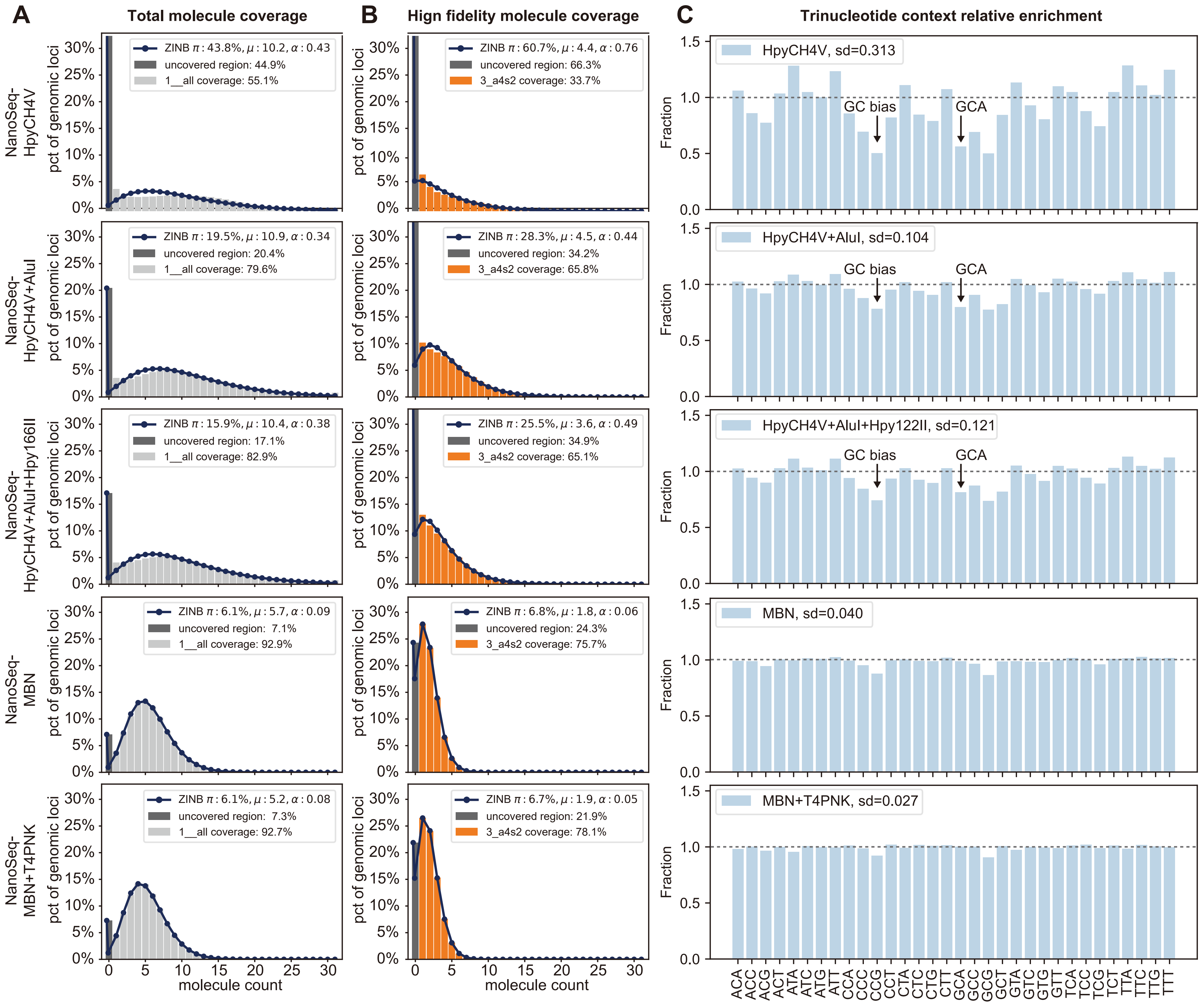
